# *Lzp* Ablation Ameliorates Dyslipidemia and Suppresses Atherosclerosis by Reducing Circulating Apolipoprotein B-Containing Lipoproteins

**DOI:** 10.1101/2025.09.24.678175

**Authors:** Kunyan He, Xin Ku, Ziyao Chen, Chenyue Wu, Chiayu Liao, Lin Li, Hongjin Qu, Peicheng Hong, Shihao Bai, Xiaofang Cui, Xianbin Su, Ze-Guang Han

## Abstract

As the principal regulator of systemic lipid homeostasis, the liver uniquely orchestrates dietary lipid assimilation, *de novo* lipogenesis, very-low-density lipoprotein (VLDL) assembly and secretion, and clearance of atherogenic lipoprotein remnants. This central role positions liver-specific molecular targets as critical therapeutic nodes for mitigating hyperlipidemia and halting atherosclerosis progression. The liver-specific protein Lzp (also named as OIT3), previously shown to stabilize apolipoprotein B (ApoB), the core structural component of triglyceride-rich lipoproteins, has an undefined role in vascular disease pathogenesis. Here, utilizing *ApoE*^−/−^mice, a well-established model that closely mimics human atherosclerosis, we demonstrate that *Lzp* deletion reduces plasma cholesterol, triglycerides, and ApoB levels under both chow and Western diets, concomitant with a marked attenuation of aortic plaque burden. Additionally, Lzp deficiency reduces hepatic and circulating ApoB levels in hyperlipidemic *ApoE*^−/−^ mice, without exacerbating hepatic steatosis or injury. Mechanistically, *Lzp* ablation is anticipated to impair hepatic VLDL-ApoB secretion based on our previous findings, resulting in reduced circulating VLDL, intermediate density lipoprotein (IDL) and low-density lipoprotein (LDL) particles, attenuated lipid deposition, and suppressed macrophage-driven plaque inflammation. These results underscore the role of Lzp as a key regulator of systemic lipid metabolism and identify it as a potential candidate for further investigation toward therapeutic intervention of atherosclerotic cardiovascular disease.

**Graphic abstract:** 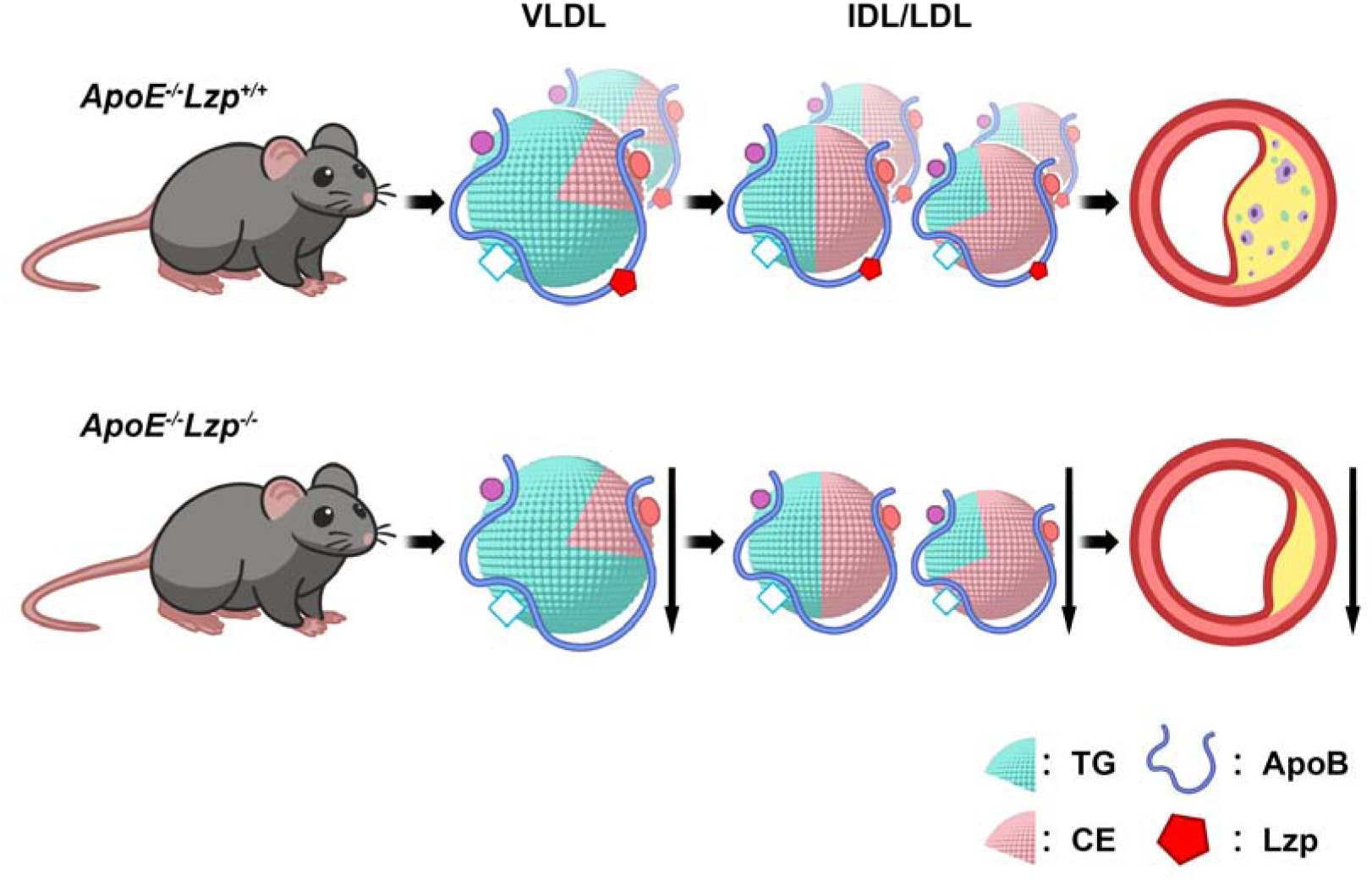

**Hightlights:**

- Lzp deficiency attenuates atherosclerosis and improves plasma lipid profiles in *ApoE*^−/−^ mice fed chow or Western diets.
- Lzp deficiency reduces hepatic and circulating apolipoprotein B levels in hyperlipidemic *ApoE*^−/−^ mice without exacerbating hepatic steatosis or injury.
- Lzp represents a candidate therapeutic target, warranting further mechanistic and translational studies for atherosclerosis.

## Introduction

Atherosclerotic cardiovascular disease (ASCVD), the global leading cause of mortality, is driven by dyslipidemia characterized by elevated low-density lipoprotein cholesterol (LDL-C), Lp(a), and triglyceride-rich lipoproteins (TGRL)^1–4^. Those atherosclerotic particles include the non-exchangeable structural scaffold apolipoprotein B (ApoB). Current lipid-lowering therapies targeting LDL-C, including statins (30-50% LDL-C lowering and 25–35% cardiovascular risk reduction) and PCSK9 inhibitors (50–60% LDL-C lowering and 15% cardiovascular risk attenuation), form the cornerstone of ASCVD management^5–7^. Despite optimal therapies, substantial residual risk persists, highlighting an urgent need for therapies targeting upstream determinants of atherogenic lipoprotein production, particularly those modulating ApoB metabolism.

As the principal orchestrator of systemic lipid homeostasis, the liver coordinates dietary lipid assimilation, *de novo* lipogenesis, VLDL assembly/secretion, coupled with hepatic clearance of atherogenic lipoprotein remnants—mechanistically interconnected processes that collectively drive atherosclerotic pathogenesis. Pharmacologic strategies increasingly focus on precision modulation of hepatic pathways. For instances, antisense oligonucleotides (ASOs), such as mipomerson, decrease ApoB translation. Small interference RNA (siRNA) therapies (inclisiran) and monoclonal antibodies (evinacumab) stabilize LDL receptor expression through PCSK9 inhibition^8–12^. Recently, emerging precision therapies have further expanded this repertoire. For example, ANGPTL3 inhibitors augment lipoprotein lipase (LPL) activity to accelerate TGRL clearance. Additionally, ASOs and siRNA selectively silence apolipoproteins C-III (ApoCIII) and Lp(a), mitigating triglyceride-driven residual risk and Lp(a) pathogenicity^13–16^. Despite these advances that herald a paradigm shift from broad lipid modulation to molecularly targeted interventions, therapeutic pleiotropy remains a critical barrier. PCSK9 inhibitors enhance LDL clearance, but their benefits are limited by a modest effect on Lp(a) reduction and a finite capacity for lowering cardiovascular risk, while the efficacy of mipomersen, which suppresses ApoB synthesis, is counterbalanced by side effects such as hepatic steatosis and injection-site reactions^12,17,18^. These limitations underscore the unmet demand for hepatic regulators of ApoB-lipoprotein production, metabolism, and clearance without disrupting systemic metabolic homeostasis or triggering compensatory pathways.

Lzp (liver-specific zona pellucida domain-containing protein), also known as OIT3, has recently been implicated in lipid metabolism^19^. Our prior work revealed that Lzp binds intracellular ApoB, shielding it from ubiquitin-proteasomal degradation, thereby gatekeeping VLDL secretion. Genetic *Lzp* ablation in mice depletes hepatic ApoB pools, reduces VLDL secretion, and lowers circulating triglycerides (TG)^19^. Despite these interesting metabolic changes, the role of Lzp in atherogenesis remains uncharacterized. Here, we interrogate the hypothesis that Lzp deficiency attenuates atherosclerosis by suppressing ApoB-containing lipoprotein secretion.

In the present work, we employed *ApoE^−/−^* mice, a well-established model of impaired lipid clearance and spontaneous atherogenesis^20^, to generate double knockout (*ApoE^−/−^ Lzp^−/−^*) mice. Our data revealed that *Lzp* deletion significantly reduced circulating cholesterol (CHOL), TG, and ApoB levels in *ApoE^−/−^* mice under both chow and Western diet conditions, concomitant with diminished atherosclerotic plaque burden. These findings establish Lzp as a previously unrecognized nexus linking hepatic ApoB regulation to systemic atherosclerosis, offering a mechanistic rationale for targeting Lzp to mitigate ASCVD risk.

## Materials & Methods

### Mice

All studies were approved by the Institutional Animal Care and Use Committee, Shanghai Jiao Tong University (Shanghai, China) and adhere to the Guide for the Care and Use of Laboratory Animals. We generated *Lzp* knockout mice (*Lzp^−/−^*) as previously described^21^. *ApoE^−/−^* mice on the C57BL/6J background were obtained from Bin Zhang’s Laboratory at Shanghai Jiao Tong University. *ApoE^−/−^* and *Lzp^−/−^* mice were crossbred to generate *ApoE^−/−^Lzp^+/+^*and *ApoE^−/−^Lzp^−/−^* littermates. The male mice were given a chow diet for 10 months or a high-cholesterol diet (0.15% cholesterol, 21% fat, Dyets, 100244) for 12 weeks starting from 6 weeks of age.

### Analysis of plasma and FPLC fractions

Plasma total cholesterol (TC) and triglyceride (TG) concentrations were quantified using enzymatic colorimetric assays (CHOD-PAP for TC, GPO-PAP for TG; Biosino Bio-technology and Science Inc.). Pooled plasma samples (20 μL per mouse; n = 6–8 mice per group) were centrifuged at 13,000g for 10 min at 4°C to remove chylomicron-rich supernatant. The samples were then subjected to fast protein liquid chromatography (FPLC) fractionation using a Superose 6 10/300 GL column (Cytiva) equilibrated with phosphate-buffered saline (PBS, pH 7.4). Elution was performed at a flow rate of 0.5 mL/min, with 0.5 mL fractions collected for subsequent analysis. Cholesterol and TG levels in each fraction were determined enzymatically using the aforementioned assays. ApoB were quantified via ELISA (Abcam, ab230932) in two compartments: (1) unfractionated plasma, and (2) FPLC peak fractions corresponding to VLDL and LDL/intermediate-density lipoprotein (IDL), as identified by cholesterol distribution profiles.

### Quantifying atherosclerosis

Following cardiac puncture, mice were transcardially perfused with 10 mL of ice-cold phosphate-buffered saline (PBS). The entire aortic tree, from the ascending aorta to the iliac bifurcation, was excised and fixed in 4% neutral-buffered formalin for 24 hours. After meticulous removal of adventitial tissue under a dissecting microscope, aortas were stained *en face* with Oil Red O (ORO; Sigma-Aldrich) to visualize lipid deposition. Stained aortas were longitudinally incised, flattened on plastic films, and imaged using a camera under standardized lighting conditions. Concurrently, hearts were harvested for aortic root analysis. Tissues were embedded in optimal cutting temperature (OCT) compound (Sakura Finetek), and cryosectioned. Serial 10-μm cross-sections spanning the aortic sinus (from the first appearance of the aortic valve leaflets to their complete disappearance, at 100-μm intervals) were collected onto adhesive slides. Atherosclerotic lesions were assessed in three complementary assays: ORO staining for lipid content; Hematoxylin & eosin (H&E) for morphology; Masson’s trichrome for fibrosis. Stained sections were imaged using an Olympus VS200 slide scanner with a 20× objective. Lesion quantification, including total vessel area, plaque area, and lumen area, was performed using ImageJ software (NIH) by investigators blinded to genotype. For immunofluorescence, cryosections were rehydrated in PBS, fixed in 4% paraformaldehyde (PFA) for 30 minutes, permeabilized with 0.1% Triton X-100/PBS for 15 minutes, and blocked with 5%BSA/PBS. Primary antibodies targeting macrophages (anti-CD68, clone KP1; Abcam ab283654; 1:200) and smooth muscle cells (anti-αSMA; Abcam ab5694; 1:500) were applied overnight at 4°C. Isotype-matched IgG (same concentrations/dilutions) served as negative controls. Sections were incubated with FITC-Conjugated secondary antibody (1:1000; Invitrogen) for 1 hour at room temperature. Images were captured using Olympus microscope.

### Liquid Chromatography-Tandem Mass Spectrometry (LC-MS/MS) methods

Plasma proteomic samples were prepared as previously described. Briefly, 2 μL of plasma was denatured at 37°C for 30 minutes with 6 M urea, reduced at 55°C for 30 minutes using 5 mM tris(2-carboxyethyl) phosphine (TCEP), alkylated with 6.25 mM iodoacetamide (RT, 1 hour in the dark), and digested with sequencing-grade trypsin (Promega; 1:20 w/w enzyme: protein) in 50 mM ammonium bicarbonate (pH 8.0) at 37°C for 16 hours. Peptides were desalted using C18 solid-phase extraction (SPE) cartridges (Thermo Fisher Scientific, 60108-301; 100 mg/1 mL), dried under vacuum, and reconstituted in 0.1% formic acid (FA). Approximately 1 μg of peptides was loaded onto an analytical column (Inspire C18; Dikma Technologies, 88305B; 150 mm × 75 μm i.d., 3 μm particle size, self-packed) equilibrated with 0.1% FA. Chromatographic separation was performed on a Dionex Ultimate 3000 nanoLC system (Thermo Fisher Scientific). Eluted peptides were analyzed using an Orbitrap Fusion mass spectrometer (Thermo Fisher Scientific) operated in data-independent acquisition (DIA) mode. Raw files were processed through DIA-NN (v1.8.1) in library-free mode against a *Mus musculus* UniProt/SwissProt database (release 2023_03; 17,052 entries). the top six fragments were utilized for peptide identification and quantification.

### Analysis of Liver

Proteins were extracted from mouse liver tissues using homogenization in cell lysis buffer, and ApoB levels were quantified with an ELISA kit. Concurrently, lipids were extracted from liver tissue using isopropanol, and triglyceride and total cholesterol levels were measured with enzymatic assays. Cryosections of liver tissue were stained with ORO and immediately scanned. For each sample, five randomly selected 1.1 × 1.1 mm fields were used for lipid area quantification with ImageJ. Additionally, paraffin-embedded liver sections were stained with H&E, scanned, and five 20× magnification fields per sample were evaluated for NAFLD activity score (NAS) according to the NASH Clinical Research Network scoring system^22^.

### Statistics

Data are presented as mean ± SEM. Statistical analyses were performed using GraphPad Prism. Normality was assessed using Shapiro-Wilk test. For data that followed a Gaussian distribution, homoscedasticity was evaluated with an F-test. Based on these assessments, between-group comparisons were made using: an unpaired Student’s t-test (for parametric data with equal variances), Welch’s corrected t-test (for parametric data with unequal variances), or a nonparametric test (Mann-Whitney U test). Statistical significance was considered at the level of *P* < 0.05.

## Results

### Lzp deficiency attenuates atherosclerosis progression in *ApoE^−/−^* mice

To investigate the role of Lzp in atherosclerosis, we generated *ApoE^−/−^Lzp^+/+^* and *ApoE^−/−^Lzp^−/−^*mice and maintained them on a chow diet for 10 months starting at 6 weeks of age. Terminal analyses revealed no significant differences in absolute body weight, liver weight, or liver-to-body weight ratio between genotypes (Figure S1), demonstrating that Lzp deficiency does not induce hepatotoxicity or visibly perturb hepatic function in *ApoE^−/−^* mice.

Notably, *en fac*e ORO staining revealed a significant reduction in atherosclerotic lesion area in whole aorta of *ApoE^−/−^Lzp^−/−^*mice compared to *ApoE^−/−^Lzp^+/+^*controls (Figure 1A-B). Cross-sectional analysis of aortic roots corroborated these findings, with *ApoE^−/−^Lzp^−/−^*mice exhibiting diminished lipid deposition (ORO staining, Figure 1B), reduced plaque area (H&E staining, Figure 1C), but unchanged collagen accumulation (Masson’s trichrome staining, Figure 1D).

**Figure 1.**
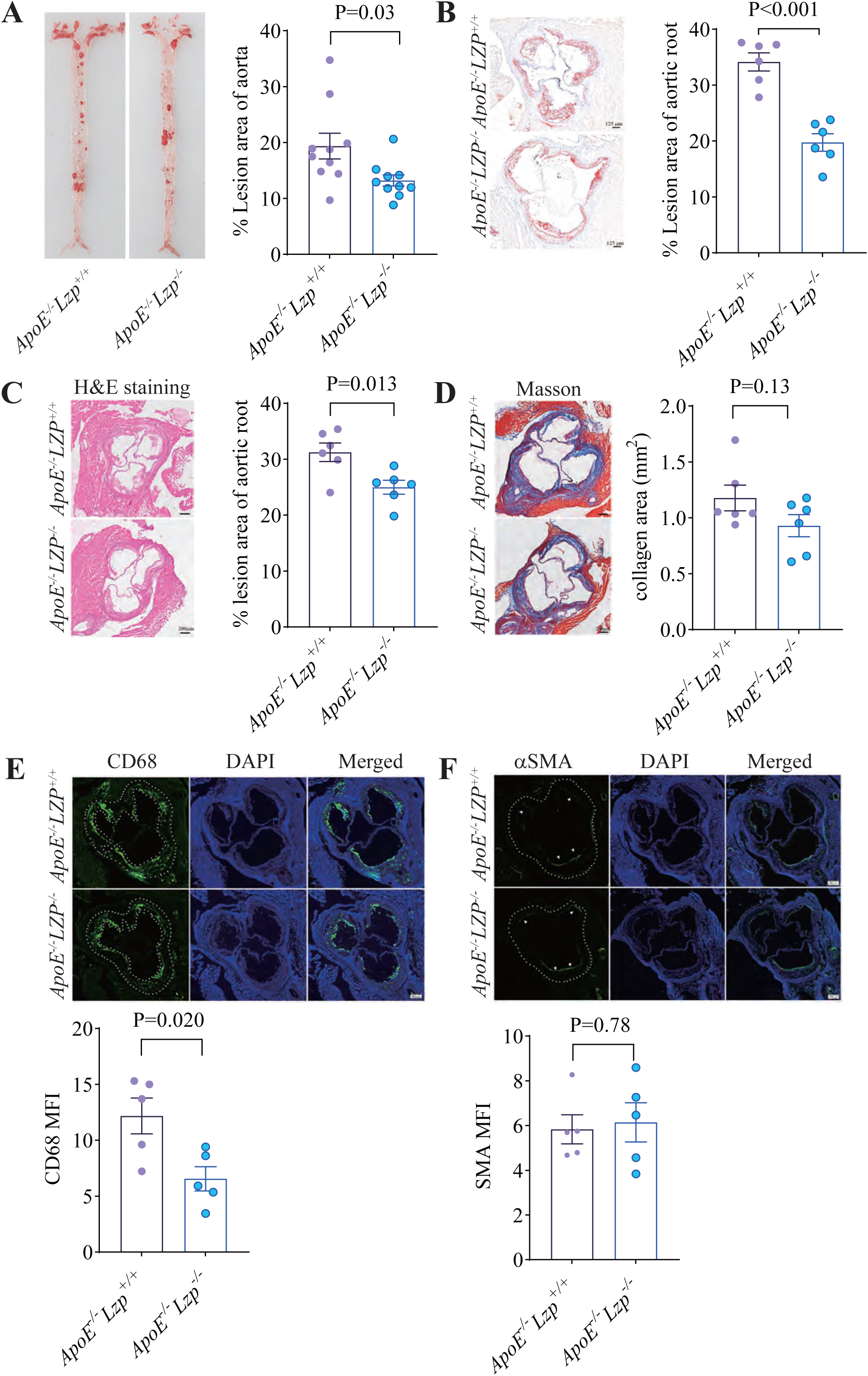
Lzp deficiency ameliorates atherosclerosis in chow diet-fed *ApoE^−/−^* mice. **A** through **F**, Six-week-old *ApoE^−/−^ Lzp^+/+^* and *ApoE^−/−^ Lzp^−/−^* mice were fed a chow diet for 10 months. **A**. Representative images (left) and quantification of oil red O (ORO)-stained *en face* aortas are shown. n = 7 for each group. **B** through **D**. Representative images (left) and quantification of hematoxylin and eosin (H&E) (**B**), ORO (**C**), and Masson (**D**) staining of aortic sinus cross sections are shown. **E** and **F**. Representative images (Upper panels) and quantification of median fluorescent intensity (MFI) for CD68-positive (**E**) and αSMA-positive cells (**F**) in aortic sinus cross sections are shown. n = 6 for each group. Data are expressed as mean ± SEM. Significant differences were determined by t-test, comparing *ApoE^−/−^ Lzp^−/−^* mice with *ApoE^−/−^ Lzp^+/+^* control. Bar lengths as indicated.

Furthermore, we performed immunofluorescence assays to quantify macrophage infiltration and smooth muscle cell (SMC) within atherosclerotic lesion area. The results revealed a significant reduction in macrophage infiltration, as indicated by a decreased CD68 cell density (Figure 1E), whereas the area occupied by α-smooth muscle actin (α-SMA) cells, a marker of SMC, remained comparable between *ApoE^−/−^Lzp^+/+^* and *ApoE^−/−^Lzp^−/−^*mice (Figure 1F). Collectively, these data suggest that Lzp deficiency attenuates atherosclerosis progression in chow diet-fed *ApoE^−/−^* mice.

### Lzp deficiency reduces plasma lipids and ApoB in *ApoE^−/−^* mice

Given our previous findings that LZP regulates ApoB-dependent VLDL secretion^19^, and the fact that *Lzp* deletion attenuates atherosclerosis progression, we comprehensively evaluated plasma lipid and apolipoprotein profiles. As expected, *ApoE^−/−^Lzp^−/−^*mice displayed pronounced reductions in plasma TG, total cholesterol (TC), and LDL-C, whereas high-density lipoprotein cholesterol (HDL-C) levels remained comparable to *ApoE^−/−^Lzp^+/+^* controls (Figure 2A-D).

**Figure 2.**
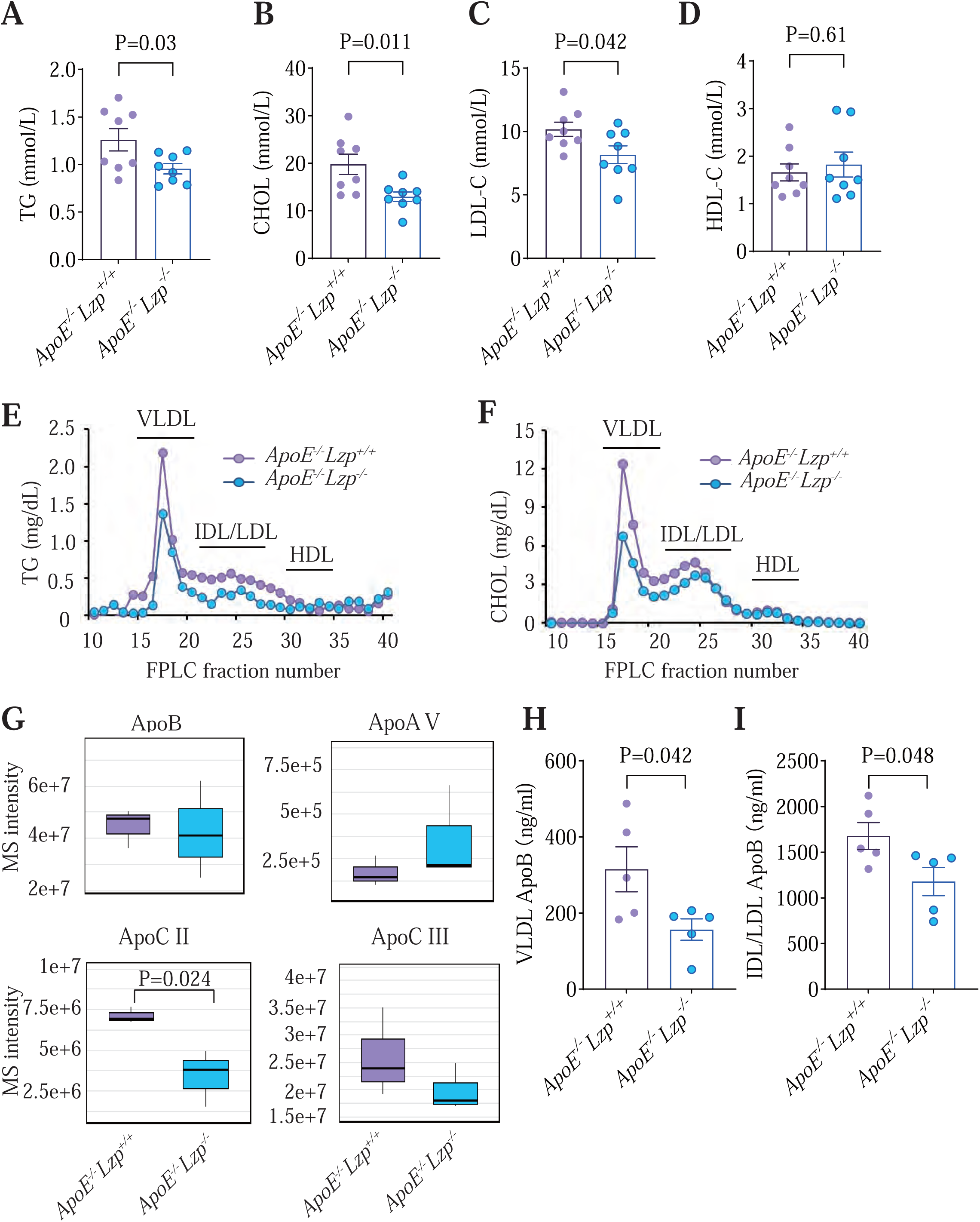
Lzp deficiency attenuates dyslipidemia in chow diet-fed *ApoE^−/−^*mice. **A** through **I**, Six-week-old *ApoE^−/−^Lzp^+/+^* and *ApoE^−/−^Lzp^−/−^* mice fed a chow diet for 10 months. Triglyceride (TG) (**A**), total cholesterol (CHOL) (**B**), low-density lipoprotein cholesterol (LDL-C) (**C**), and high-density lipoprotein cholesterol (HDL-C) (**D**) levels in plasma were quantified. **E** and **F**, plasma was pooled from each group (n = 6) and separated by fast protein liquid chromatography (FPLC). The TG and cholesterol in the fractions were determined after separation. Plasma TG (**E**) and cholesterol (CHO) (**F**) distribution across these fractions are present. **(G)** Plasma lipoproteins were quantified by data-independent acquisition mass spectrometry (DIA-MS) (n = 3 per group). Boxplots show the quantification of ApoB, ApoA-V, ApoC-II, and ApoC-III in *ApoE^−/−^Lzp^+/+^* and *ApoE^−/−^Lzp^−/−^*mice. **H** and **I**, the ApoB levels in the VLDL (**H**) and IDL/LDL (**I)** fraction peaks obtained from fast protein liquid chromatography (FPLC) separation of plasma for cholesterol plasma were measured with an ELISA assay. Data were statistically analyzed by t-test, and the values are expressed as mean ± SEM.

To accurately resolve this lipid alterations, we employed fast protein liquid chromatography (FPLC) to fractionate lipoprotein subclasses. Interestingly, both TG and TC contents were markedly diminished in VLDL and its catabolic derivatives IDL/LDL particles, while these contents in HDL particles remained unaffected (Figure 2E-F). These data indicated that LZP deficiency specifically impairs the metabolism of ApoB containing lipoproteins.

To assess the global impact of LZP deficiency on the lipoprotein landscape, we conducted a system-wide quantitative mass spectrometry analysis of apolipoproteins. The known pro-atherogenic apolipoproteins ApoB and ApoCIII, the core components of VLDL particles, were markedly reduced in plasma of *ApoE^−/−^Lzp^−/−^* mice. While the LPL co-activator ApoCII was diminished, ApoAV, which enhances TG hydrolysis, was upregulated in these mice (Figure 2G). Interestingly, other apolipoproteins associated with IDL/LDL and HDL (e.g., ApoA-I, ApoA-II, ApoA IV, ApoM, ApoCI, ApoC IV, and ApoF) were also suppressed, whereas ApoF and ApoD remained unchanged (Figure S2). To confirm these observations, we employed ELISA assay to quantify ApoB levels in isolated VLDL and LDL fractions. Consistent with the proteomic data, *ApoE^−/−^Lzp^−/−^*mice exhibited pronounced reductions in ApoB content within VLDL and LDL particles compared to *ApoE^−/−^Lzp^+/+^*mice (Figure 2H and 2I).

These results indicate that Lzp deficiency alleviates atherogenic lipoprotein burden primarily through suppression of ApoB-containing particles (VLDL, IDL and LDL), consistent with its role as a hepatic ApoB stabilizer reported in our prior work^19^. Through its dual modulation of lipoprotein biogenesis and metabolism, Lzp emerges as a pivotal regulator of lipid metabolic flux and atherosclerotic progression.

### Lzp deficiency lowers hepatic ApoB without exacerbating lipid accumulation or injury in *ApoE^−/−^* mice

Previous study reported that Lzp deficiency reduces ApoB levels and promotes lipid accumulation without overt liver damage^19^. To investigate this under hyperlipidemic conditions, we employed Lzp and ApoE double-knockout mice. As expected, hepatic ApoB protein levels were significantly reduced in *ApoE^−/−^Lzp^−/−^*mice compared to *ApoE^−/−^Lzp^+/+^* controls (Figure 3A). We next assessed hepatic lipid content and markers of liver injury. TG content was significantly reduced in the livers of *ApoE^−/−^Lzp^−/−^* mice, while total cholesterol (TC) level remained unchanged (Figure 3B and 3C). ORO staining indicated comparable lipid accumulation between genotypes (Figure 3D). Histological evaluation via H&E staining and subsequent NAFLD activity score (NAS) analysis based on the NASH Clinical Research Network scoring system revealed no signs of exacerbated steatohepatitis (Figure 3E).

**Figure 3.**
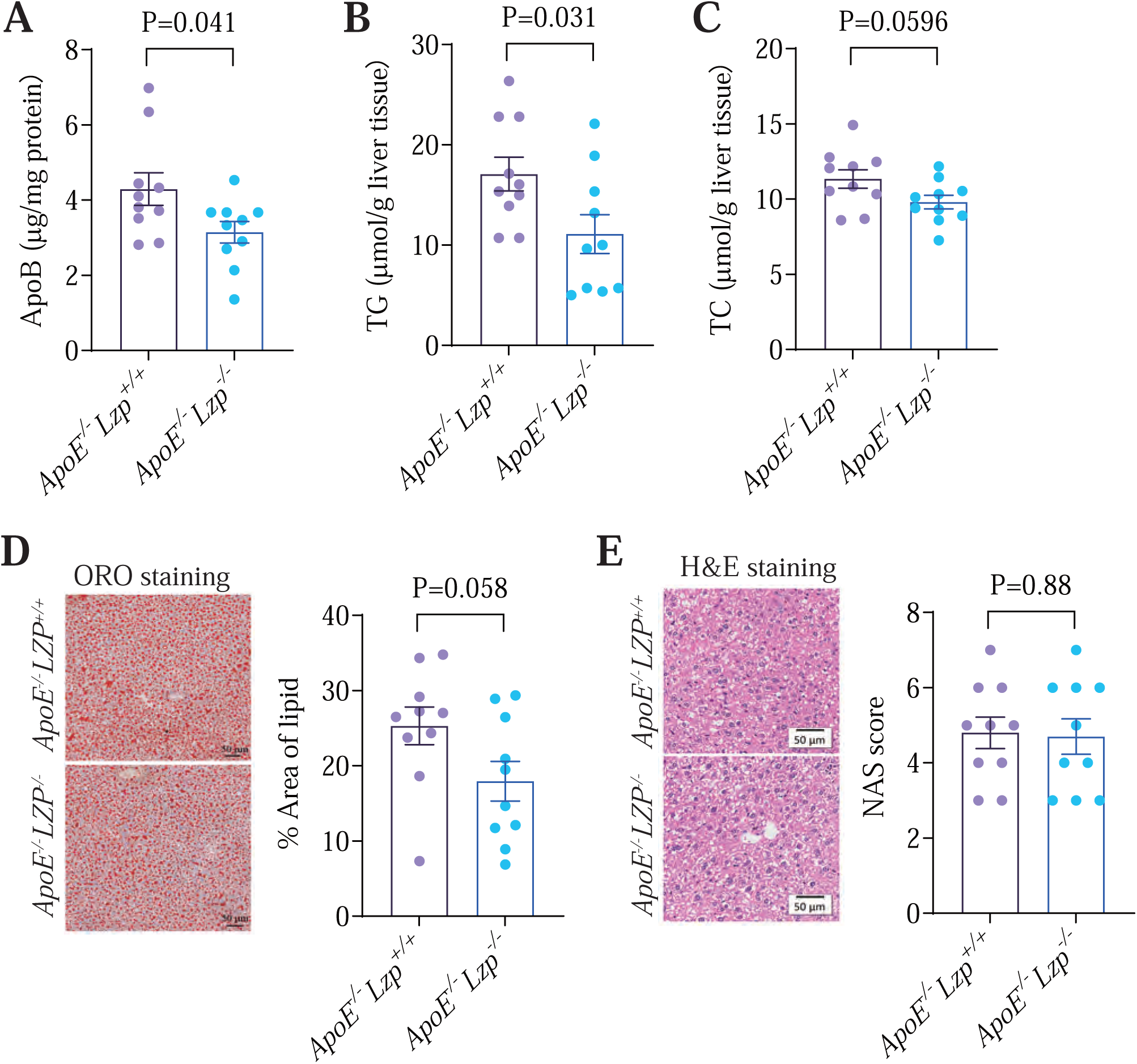
Lzp deficiency lowers hepatic ApoB without inducing lipid accumulation or injury in *ApoE^−/−^*mice. **A** through **E**, Six-week-old *ApoE^−/−^Lzp^+/+^* and *ApoE^−/−^Lzp^−/−^* mice fed a chow diet for 10 months. **(A)**. The ApoB levels in the liver tissues were measured with an ELISA assay. Triglyceride (TG) (**B**) and total cholesterol (CHOL) (**C**) of livers were quantified. Representative images (left) and quantification of ORO (**D**) and hematoxylin and eosin (H&E) (**E**) staining of liver sections are shown. Data were statistically analyzed by t-test, and the values are expressed as mean ± SEM.

qPCR analysis also showed no significant differences in the expression of genes involved in lipoprotein assembly, catabolism and uptake, lipogenesis, bile acid metabolism, or inflammation between genotypes, despite complete ablation of *Lzp* expression (Figure S3).

Together, these findings indicate that Lzp deficiency reduces hepatic ApoB and TG content without inducing additional lipid accumulation, injury, or inflammatory activation in an *ApoE^−/−^* mice.

### Lzp deficiency protects against Western diet-accelerated atherosclerosis

To further validate the atheroprotective effect of Lzp deficiency, *ApoE^−/−^Lzp^+/+^ and ApoE^−/−^Lzp^−/−^* mice were challenged with a Western diet for 12 weeks starting at 6 weeks of age. *ApoE^−/−^Lzp^−/−^*and *ApoE^−/−^Lzp^+/+^* mice exhibited comparable absolute body weight, liver weight and liver-to-body weight ratio (Figure S4A-C). Key indicators of hepatocellular integrity and functions, including plasma alanine aminotransferase (ALT), aspartate aminotransferase (AST), albumin, and glucose levels, remained unaltered (Figure S4D-G), confirming that LZP deficiency does not induce overt hepatotoxicity or significantly perturb systemic metabolic homeostasis, even under Western diet challenges.

Significantly, compared to *ApoE^−/−^Lzp^+/+^*controls, *ApoE^−/−^Lzp^−/−^* mice exhibited attenuated severity of atherosclerosis, as evidenced by reduced *en face* aortic lesion area (ORO staining, Figure 4A), diminished lipid accumulation in aortic roots (ORO staining, Figure 4B), and smaller plaque area (H&E staining, Figure 4C). In addition, plaque stability parameters were improved, with obviously decreased collagen deposition (Masson’s trichrome staining, Figure 4D) and markedly reduced intraplaque macrophages (quantified by CD68 fluorescence intensity, Figure 4E), while smooth muscle cell content remained unchanged (quantified by α-SMA fluorescence intensity, Figure 4F).

**Figure 4.**
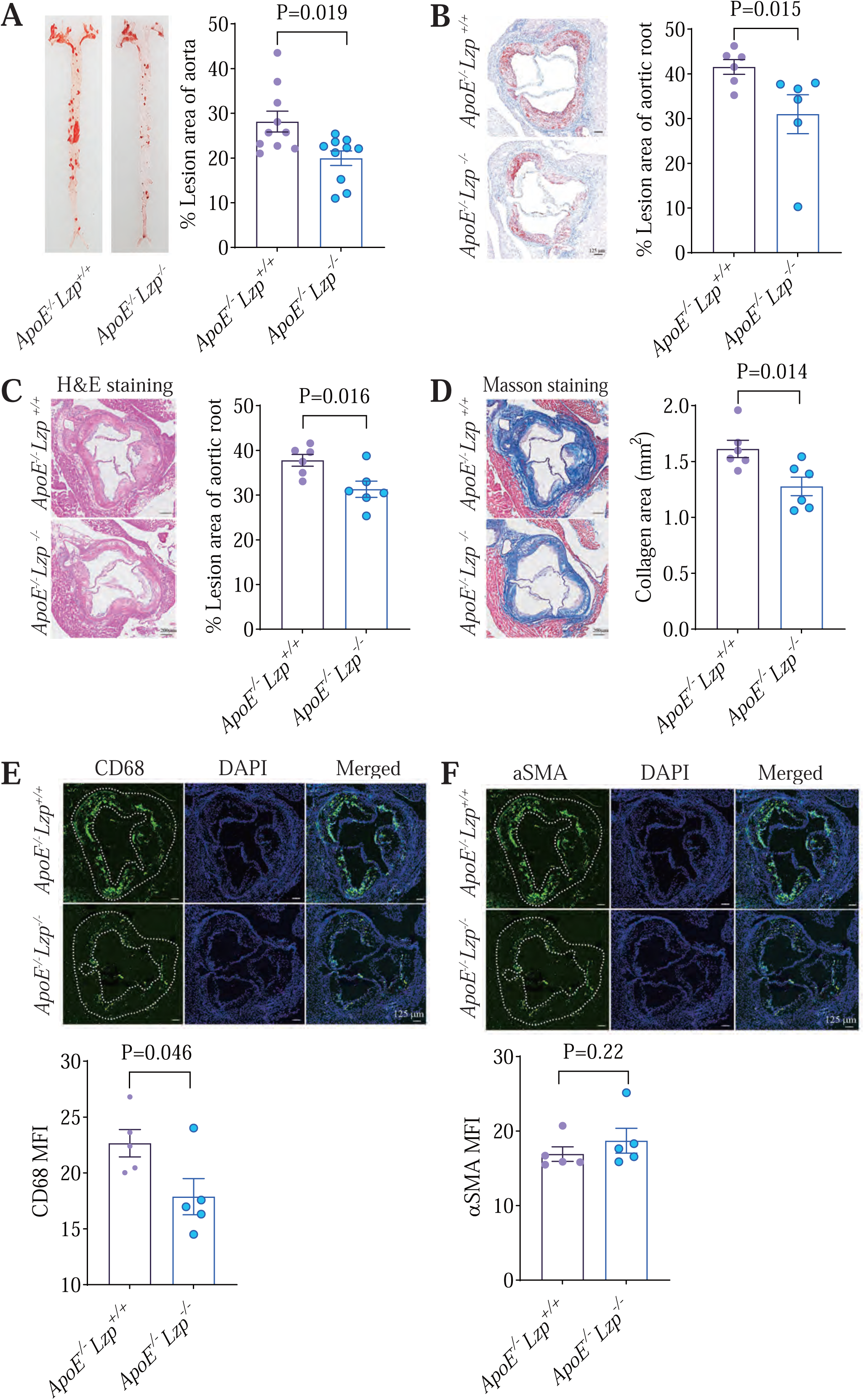
Lzp deficiency ameliorates Western diet (WD)-accelerated atherosclerosis in *ApoE^−/−^*mice. **A** through **F**, Six-week-old *ApoE^−/−^Lzp^+/+^*and *ApoE^−/−^Lzp^−/−^* mice were fed a WD for 12 weeks. **A**. Representative images (left) and quantification of ORO-stained *en face* aortas are shown. n = 10 for each group. **B** through **D**. Representative images (left) and quantification of H&E (**B**), ORO (**C**), and Masson (**D**) staining of aortic sinus cross sections were present. **E** and **F**, Representative images (Upper panels) and quantification of median fluorescent intensity (MFI) for CD68-positive (**E**) and αSMA-positive cells (**F**) in aortic sinus cross sections were shown. n = 6 for each group. Data were expressed as mean ± SEM. Significant differences were determined by Student’s t-test, comparing *ApoE^−/−^ Lzp^−/−^* mice with *ApoE^−/−^ Lzp^+/+^* control. Bar lengths as indicated.

These data confirm that Lzp deficiency confers robust atheroprotection under both chow and Wester diet conditions, supporting a diet-independent role in mitigating vascular pathology.

### Lzp deficiency ameliorates Western diet-induced dyslipidemia

Consistent with results in chow-fed mice (Figure 2), Western diet-fed *ApoE^−/−^Lzp^−/−^* exhibited improved lipid homeostasis, characterized by significantly reduced plasma TG and TC levels (Figure 5A and 5B). However, no significant changes were observed in plasma LDL-C and HDL-C levels (Figure 5C and 5D).

**Figure 5.**
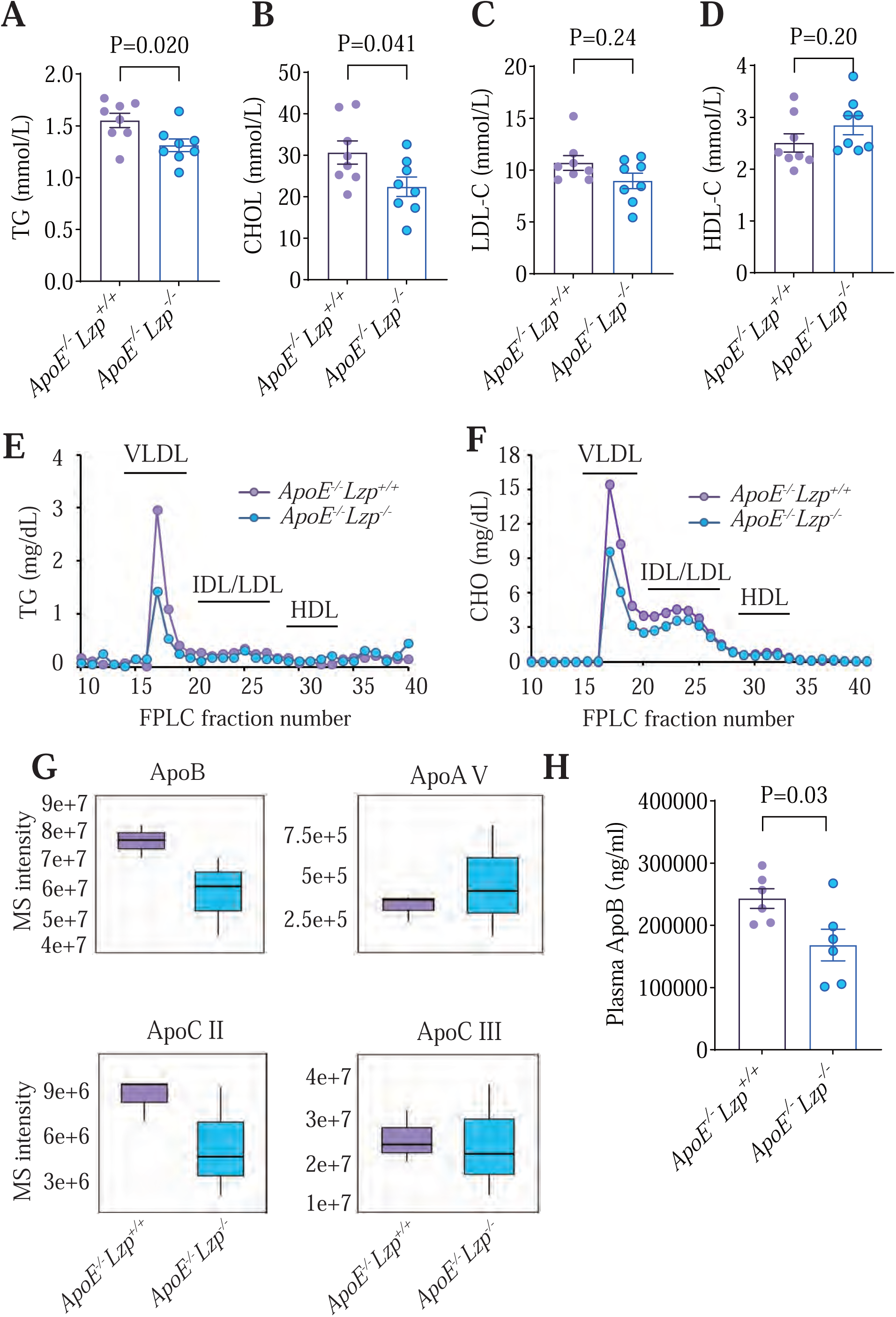
Lzp deficiency alleviates dyslipidemia in WD-fed *ApoE^−/−^* mice. **A** through **H**, Six-week-old *ApoE^−/−^Lzp^+/+^* and *ApoE^−/−^Lzp^−/−^*mice fed a WD for 12 weeks. TG (**A**), TC (**B**), LDL-C (**C**), and HDL-C (**D**) levels in plasma were quantified. **E** and **F**, plasma was pooled from each group (n = 8) and separated by FPLC. The TG and cholesterol in the fractions were determined after separation. Plasma TG (**E**) and cholesterol (CHOL) (**F**) distribution across these fractions are present. **G**. Plasma lipoproteins were quantified by DIA-MS (n = 3 per group). Boxplots show the quantification of ApoB, ApoA-V, ApoC-II, and ApoC-III between *ApoE^−/−^Lzp^+/+^* and *ApoE^−/−^Lzp^−/−^*mice. **H**. The ApoB levels in the plasma were measured with an ELISA assay. Data were statistically analyzed by t-test, and the values are expressed as mean ± SEM.

Subsequently, to accurately quantify TG and cholesterol levels across lipoprotein fractions, we also performed FPLC profiling on mouse plasma. The analysis revealed marked decreased in both TG and cholesterol within the VLDL and IDL/LDL fractions (Figure 5E and 5F), whereas levels in the HDL fraction remained unaltered.

Quantitative mass spectrometry analysis of apolipoproteins revealed selective decreases in ApoB and ApoC-II/III levels and an elevation in ApoA-V levels (Figure 5G). Also, apolipoproteins associated with IDL/LDL and HDL (ApoA-I, ApoA-II, ApoM, ApoCI, ApoC IV, and ApoF) were suppressed, whereas ApoA IV and ApoD were elevated (Figure S5). These findings were consistent with the observations in mice fed a chow diet (Figures 3 and S2). Importantly, ELISA quantification confirmed significant reductions of plasma ApoB (Figure 5H), aligning with FPLC-based lipoprotein distribution and proteomic data.

### Lzp deficiency reduces hepatic ApoB without aggravating lipid accumulation or injury in Western diet-fed *ApoE^−/−^* mice

ELISA analysis of liver tissue confirmed that Lzp deficiency significantly decreased ApoB protein levels in *ApoE^−/−^* mice fed a Western diet, supporting a role for LZP in ApoB stabilization (Figure 6A). Hepatic TG and TC levels were comparable between *ApoE^−/−^Lzp^+/+^*and *ApoE^−/−^Lzp^−/−^* mice (Figure 6B and 6C), as was lipid accumulation by ORO staining of liver sections (Figure 6D). Furthermore, NAFLD activity scores (NAS) revealed no differences between two genotypes (Figure 6E).

**Figure 6.**
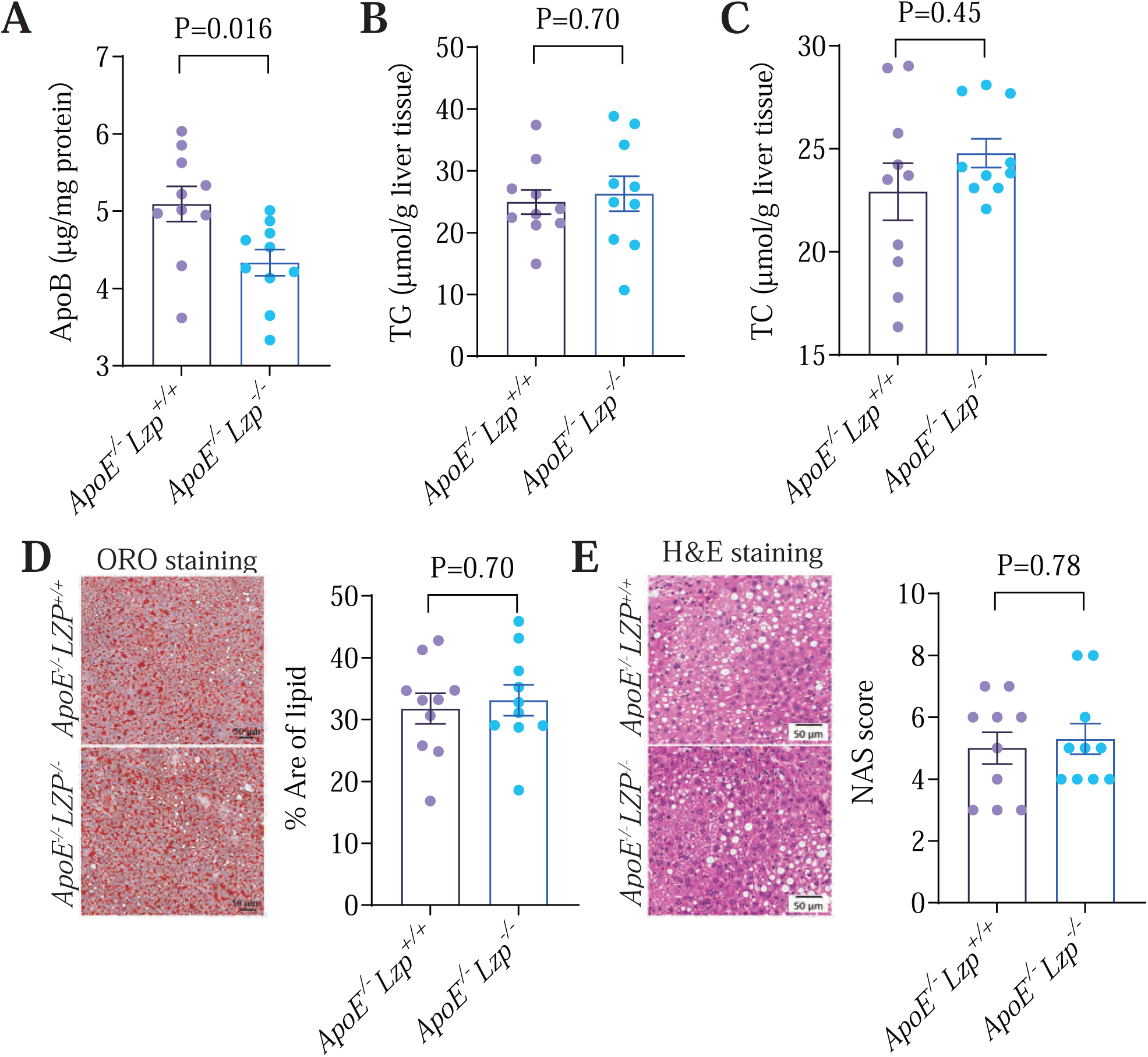
Lzp deficiency lowers hepatic ApoB without inducing lipid accumulation or injury in WD-fed *ApoE^−/−^* mice. **A** through **E**, Six-week-old *ApoE^−/−^Lzp^+/+^*and *ApoE^−/−^Lzp^−/−^* mice fed a WD for 12 weeks. **(A)**. The ApoB levels in the liver tissues were measured with an ELISA assay. Triglyceride (TG) (**B**) and total cholesterol (CHOL) (**C**) of livers were quantified. Representative images (left) and quantification of ORO (**D**) and hematoxylin and eosin (H&E) (**E**) staining of liver sections are shown. Data were statistically analyzed by t-test, and the values are expressed as mean ± SEM.

qPCR analysis also revealed no changes in expression of genes related to lipoprotein assembly, catabolism and uptake, lipogenesis, bile acid metabolism, or hepatic inflammation, except *Lzp* mRNA expression was ablated in *ApoE^−/−^Lzp^−/−^*mice (Figure S6).

Collectively, these results demonstrate that Lzp deficiency attenuates atherogenesis largely through reducing TG and cholesterol in VLDL and their metabolic products (IDL/LDL), as well as key apolipoproteins, particularly ApoB, without eliciting adverse hepatic effects.

## Discussion

Our previous study establishes LZP as a liver-specific regulator of ApoB stability and lipoprotein secretion. Building on this foundation, the current work demonstrates that global *Lzp* deletion ameliorates hyperlipidemia and attenuates atherosclerosis in *ApoE^−/−^*mice. The observed reductions in plasma cholesterol, triglycerides, and atherogenic ApoB-containing lipoproteins under both dietary regimens align with impaired hepatic VLDL-ApoB secretion, a mechanism established in our earlier findings. Together, these results suggest that LZP merits further investigation as a candidate therapeutic target for atherosclerotic cardiovascular disease.

The liver plays a central role in systemic lipid homeostasis, coordinating dietary lipid processing, *de novo* lipogenesis, lipoprotein assembly and secretion, and hepatic clearance of atherogenic remnants, making it a critical focus for therapeutic interventions against metabolic disorders such as atherosclerosis^23^. Existing approaches, including statins (HMG-CoA reductase inhibitors), fibrates (PPAR-α agonists), and apolipoprotein-directed agents like ApoB translation suppressor mipomersen, highlight the importance of hepatic targeting in lipid management^24^. Emerging evidence further implicates liver-specific proteins like CREBH (involved in remnant lipoprotein clearance) and SMLR1 (implicated in VLDL trafficking) in modulating atherogenic risk^25,26^. In line with these insights, we show that genetic ablation of Lzp reduces ApoB-enriched atherogenic particles (VLDL/LDL) and attenuates plaque burden in *ApoE^−/−^* mice, supporting a role for LZP in atherogenesis through regulation of ApoB stability and VLDL secretion. Our study delineates LZP as a previously unrecognized hepatic regulator of atherogenesis.

In this study based on *ApoE^−/−^* mouse models, quantitative proteomic profiling revealed Lzp deficiency not only reduced proatherogenic ApoB and ApoCIII (an LPL inhibitor and NLRP3 inflammasome activator) but also modulated protective apolipoproteins such as ApoA-V. It is notable that reduced ApoC-II (an LPL coactivator) and ApoA-I (a major HDL component) were also detected (Figures 2, 5, S2 and S5). The overall atheroprotective phenotype implies that the benefits of reducing ApoB-containing lipoproteins outweigh potential drawbacks from modulating other apolipoproteins; nevertheless, the full physiological implications of these changes require further study.

Interestingly, using well-established *ApoE^−/−^* mouse models^20,27^, our dietary comparative analyses revealed nuanced therapeutic dynamics. Atherosclerosis was attenuated under both chow and Western diet conditions. However, the reduction in collagen deposition, as evaluated by Masson’s trichrome staining, was less pronounced in mice fed a chow diet (Figures 1 and 4). This observation suggests that dietary and age influences on plaque composition and remodeling (Figures 1 and 4).

The reduced collagen deposition in plaques of Western diet-fed *ApoE^−/−^ Lzp^−/−^* mice implies that Lzp may exert context-dependent effects on plaque stability, potentially exacerbating early-stage fibrotic remodeling while attenuating late-stage calcification (Figure 4D). In early atherogenesis (Figure 1D, chow model), reduced ApoB availability likely limits lipoprotein retention and subendothelial modification. Conversely, in advanced lesions (Western diet), modulation of ApoC-III (an NLRP3 inflammasome activator) and collagen remodeling may additionally influence inflammatory and fibrotic pathways. This context-dependence mirrors clinical observations where lipid-lowering efficacy varies with disease stage, underscoring the need for patient-stratified therapeutic approaches.

This work positions LZP within the expanding landscape of hepatic targets for precision atherosclerosis therapy. The liver-specific expression of LZP makes it amenable to silencing via GalNAc-conjugated siRNAs or CRISPR-based epigenetic editing, approaches that may offer improved specificity over small molecule inhibitors. Furthermore, as an upstream regulator of ApoB, LZP inhibition may act synergistically with existing LDL-C-lowering agents (e.g., statins, bempedoic acid) to address residual cardiovascular risk.

Several limitations of this study should be acknowledged. The use of a global knockout model leaves open the possibility of extrahepatic contributions, for example, from macrophages or other tissues expressing LZP. In addition, direct measurement of VLDL-ApoB secretion in the ApoE / background, evaluation of sex-specific effects, and validation using hepatocyte-specific *Lzp*-knockout models will be essential to firmly establish LZP as a translational target. Future studies should also clarify LZP’s mechanistic roles in ApoB vesicular trafficking and post-translational regulation, the temporal effects of LZP modulation on plaque progression and regression, and the efficacy of pharmacological LZP inhibition in humanized models.

## Author contributions

K-Y. He: Investigation, formal analysis, visualization, methodology, writing-original draft, funding acquisition. X. Ku: Investigation, data curation, methodology. Z-Y. Chen: Methodology. C-Y. Wu: Methodology. J-Y.Liao: Methodology. H-J. Qu: Resources, methodology. S-H. Bai: Methodology. P-C. Hong: Methodology. X-B. Su: Resources, data curation. X-F. Cui: Resources, data curation. Z-G. Han: Conceptualization, data curation, supervision, writing-review, funding acquisition.

## Data availability statement

All data presented in the manuscript are available from the corresponding author upon reasonable request.

