## Supplementary material for "*Lzp* Ablation Ameliorates Dyslipidemia and Suppresses Atherosclerosis by Reducing Circulating Apolipoprotein B-Containing Lipoproteins": Figure S1-S6

**Supplementary Figure legends**

**Figure S1. The analysis of absolute body weight, liver weight, or liver-to-body weight ratio in chow diet-fed *ApoE^−/−^* mice.** The body weight (A), liver weight (B), and liver-to-body ratio (C) were measured in *ApoE^−/−^Lzp^+/+^* and *ApoE^−/−^Lzp^−/−^* mice (n = 7 per group). Data were statistically analyzed by t-test, and the values are expressed as mean ± SEM.

**Figure S2. Quantification of plasma lipoproteins in chow diet-fed *ApoE^−/−^* mice.** Plasma lipoproteins were quantified by data-independent acquisition mass spectrometry (DIA-MS). Boxplots show the quantification of these lipoproteins as indicated in *ApoE^−/−^Lzp^+/+^* and *ApoE^−/−^Lzp^−/−^* mice (n = 3 per group), as indicated.

**Figure S3. Quantification of genes’ expression associated hepatic lipoprotein assembly, catabolism and uptake, lipogenesis, bile acid metabolism, or inflammation in chow diet-fed *ApoE^−/−^* mice.** Quantification of genes indicated in liver tissues of *ApoE^−/−^Lzp^+/+^* and *ApoE^−/−^Lzp^−/−^* mice (n = 10). Data were statistically analyzed by t-test, and the values are expressed as mean ± SEM.

**Figure S4. The analysis of absolute body weight, liver weight, liver-to-body weight ratio, metabolic and hepatic functions in Western diet-fed *ApoE^−/−^* mice.** The body weight (A), liver weight (B), liver-to-body ratio (C), the levels of blood alanine aminotransferase (ALT) (D), aspartate aminotransferase (AST) (E), albumin (ALB) (F) and glucose (GLU) (G) were measured in *ApoE^−/−^Lzp^+/+^* and *ApoE^−/−^Lzp^−/−^* mice (n = 8 or 10 per group). Data were statistically analyzed by t-test, and the values are expressed as mean ± SEM.

**Figure S5. Quantification of plasma lipoproteins in Western diet-fed *ApoE^−/−^* mice.** Plasma lipoproteins were quantified by data-independent acquisition mass spectrometry (DIA-MS). Boxplots show the quantification of these lipoproteins as indicated in *ApoE^−/−^Lzp^+/+^* and *ApoE^−/−^Lzp^−/−^* mice (n = 3 per group), as indicated.

**Figure S6. Quantification of genes’ expression associated hepatic lipoprotein assembly, catabolism and uptake, lipogenesis, bile acid metabolism, or inflammation in Western diet-fed *ApoE^−/−^* mice.** Quantification of genes indicated in liver tissues of *ApoE^−/−^Lzp^+/+^* and *ApoE^−/−^Lzp^−/−^* mice (n = 10). Data were statistically analyzed by t-test, and the values are expressed as mean ± SEM.
