## Supplementary figures and images for "*Lzp* Ablation Ameliorates Dyslipidemia and Suppresses Atherosclerosis by Reducing Circulating Apolipoprotein B-Containing Lipoproteins"

### Supplementary Figure legends

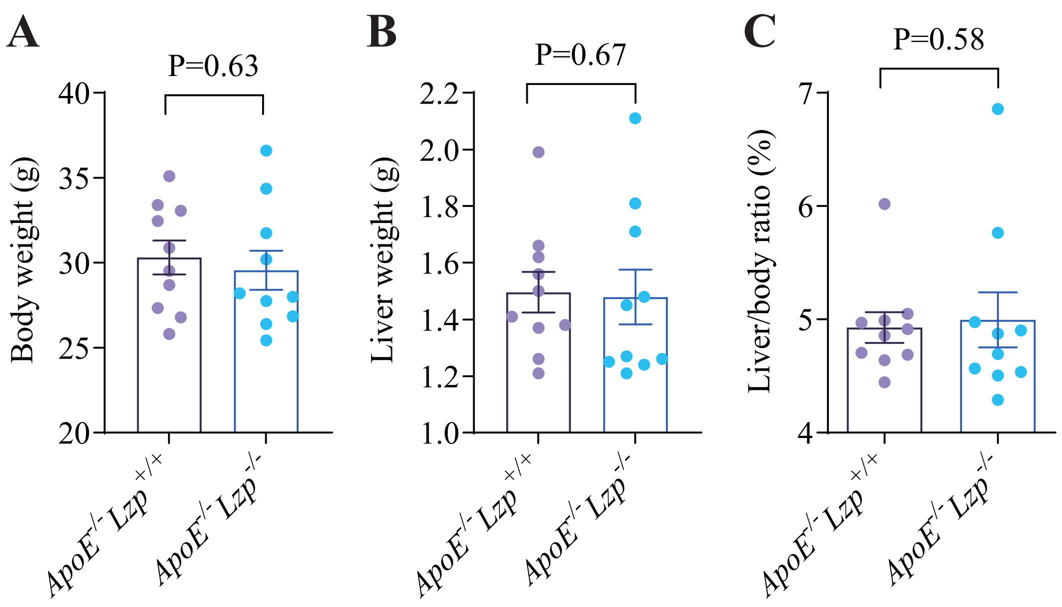

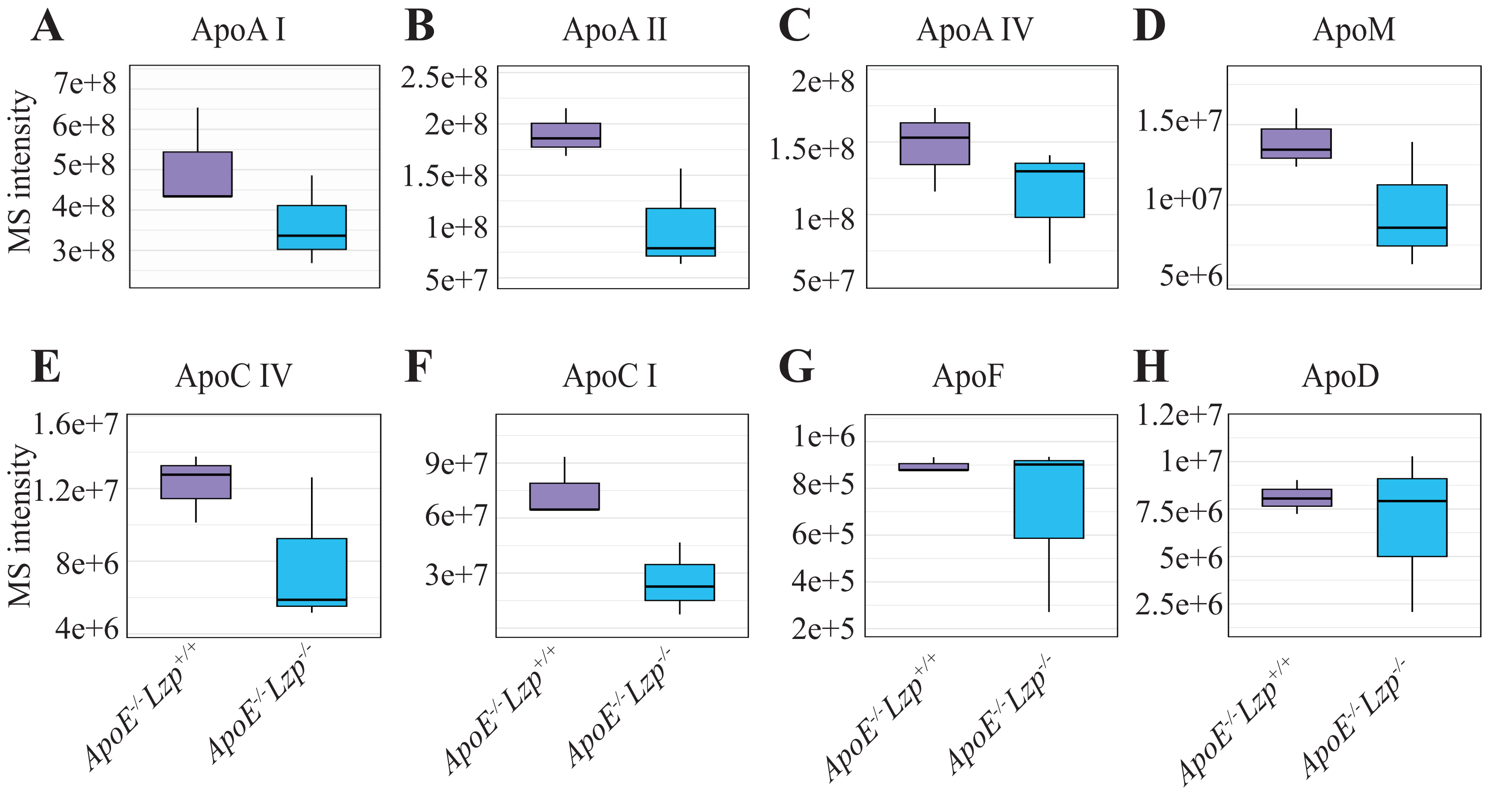

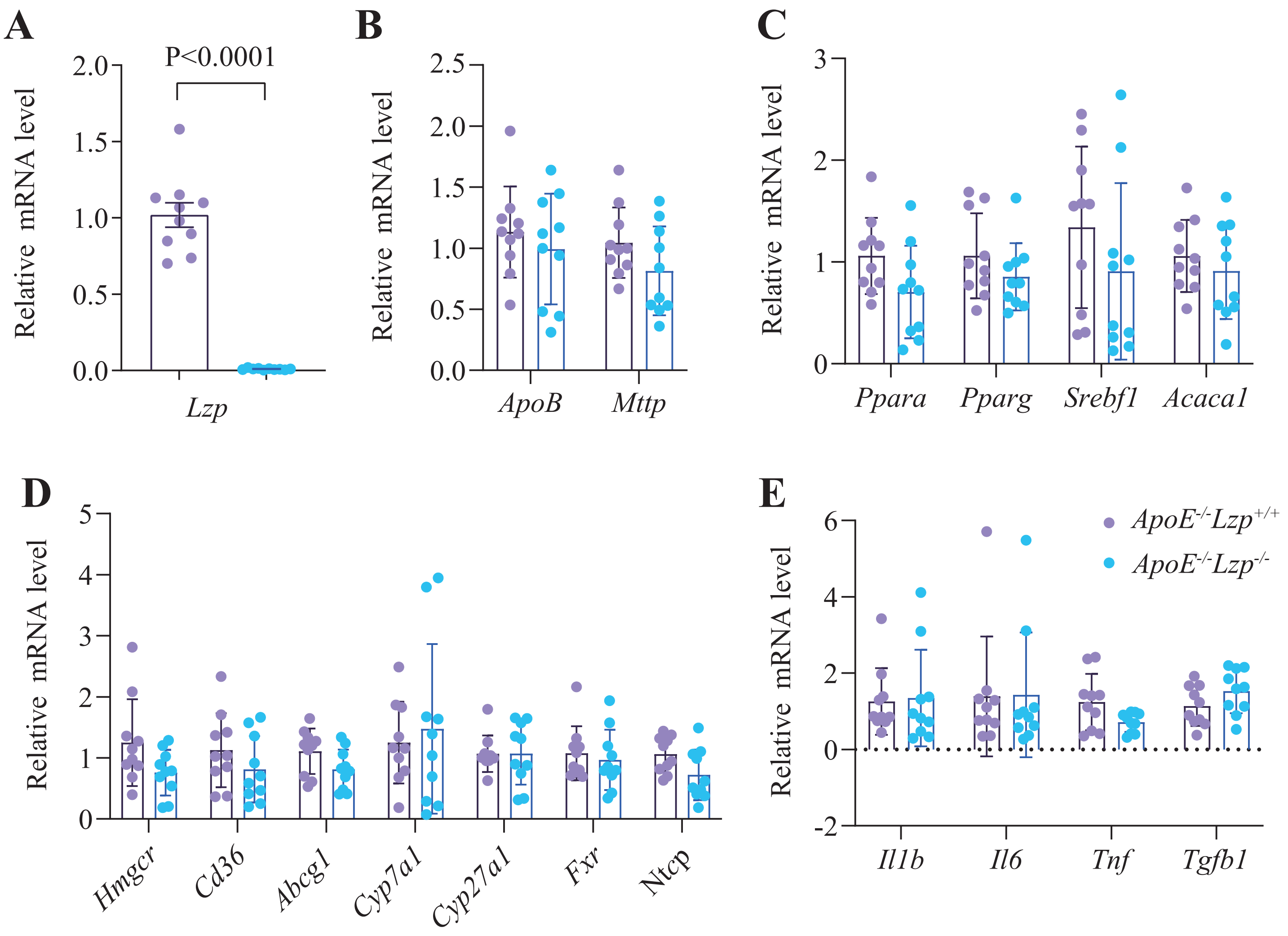

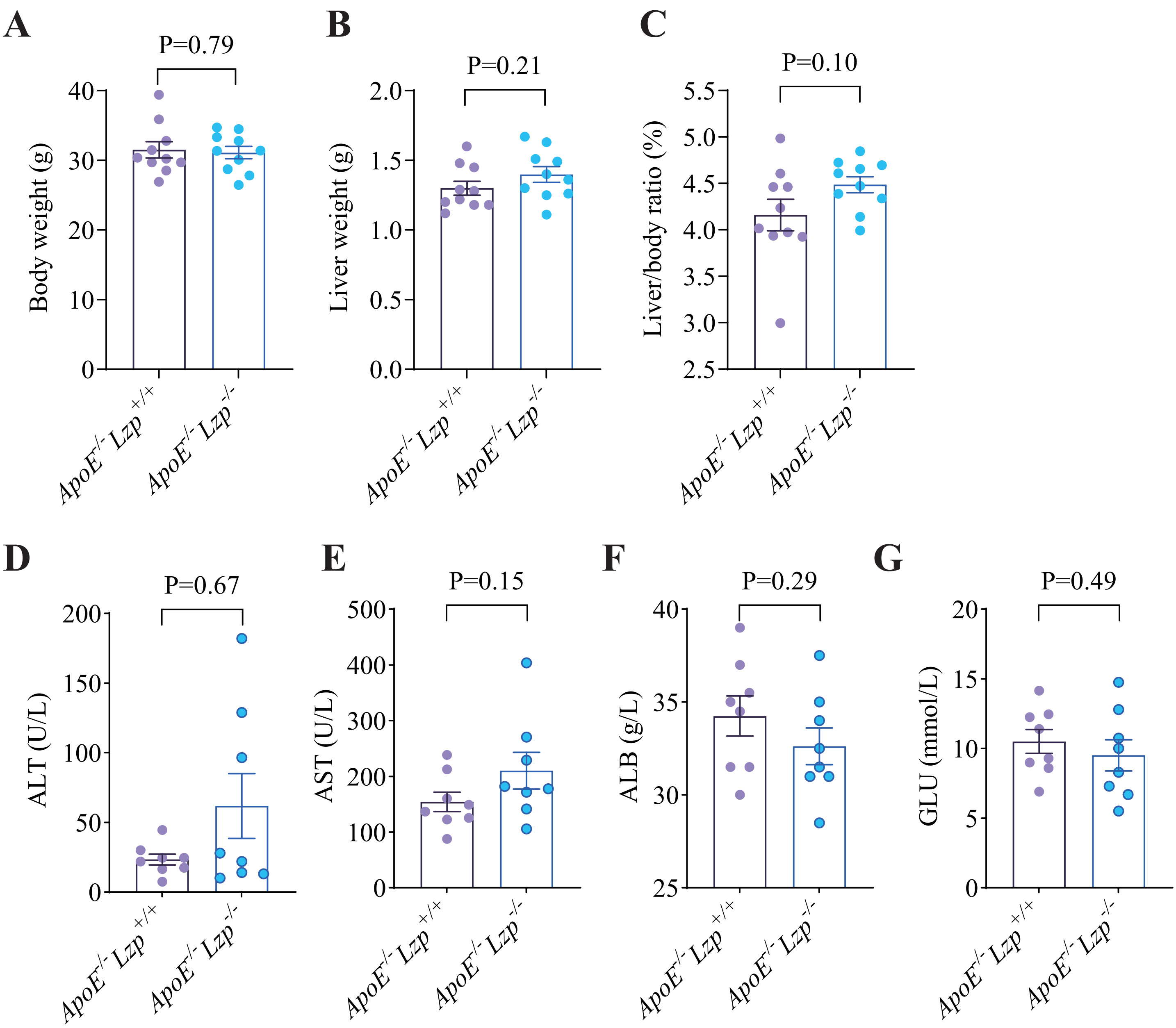

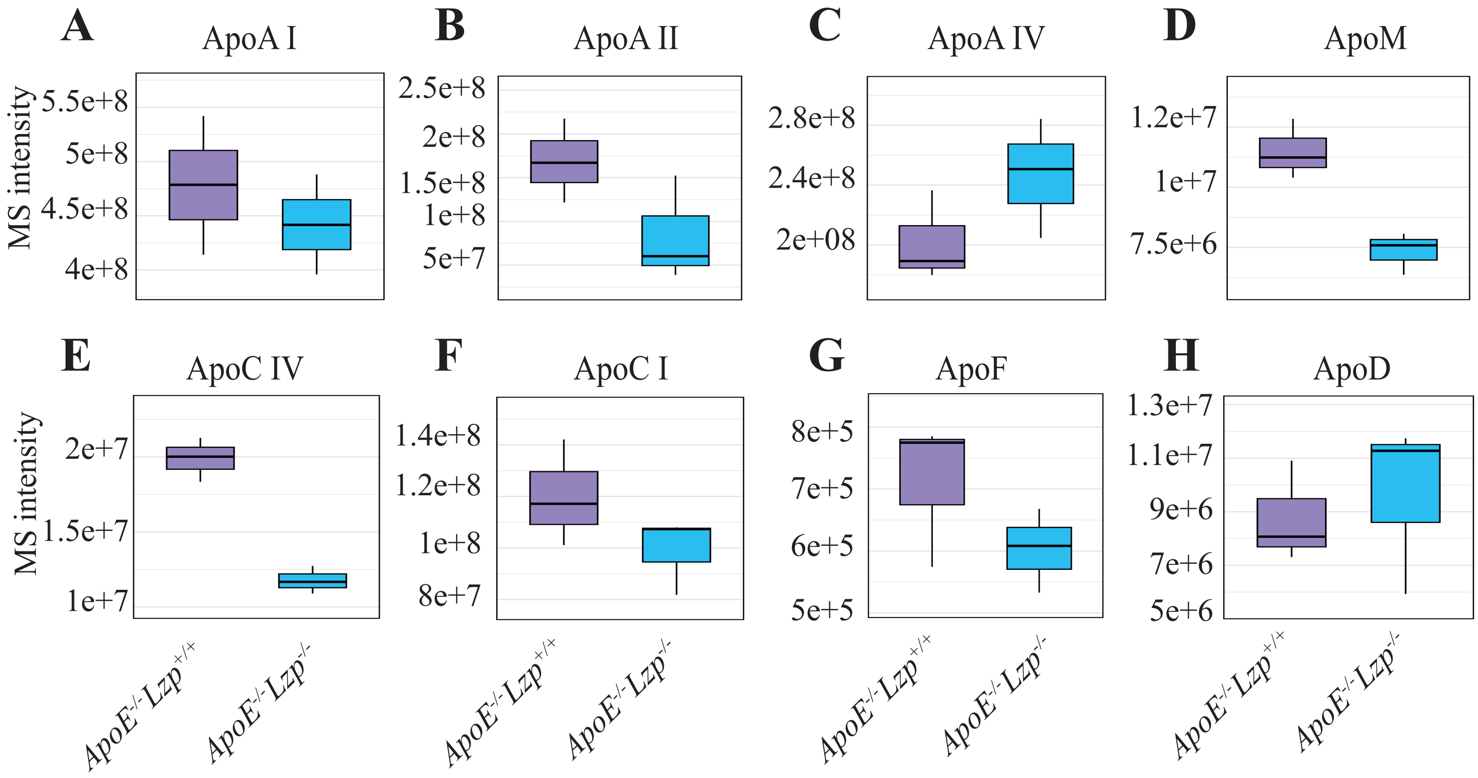

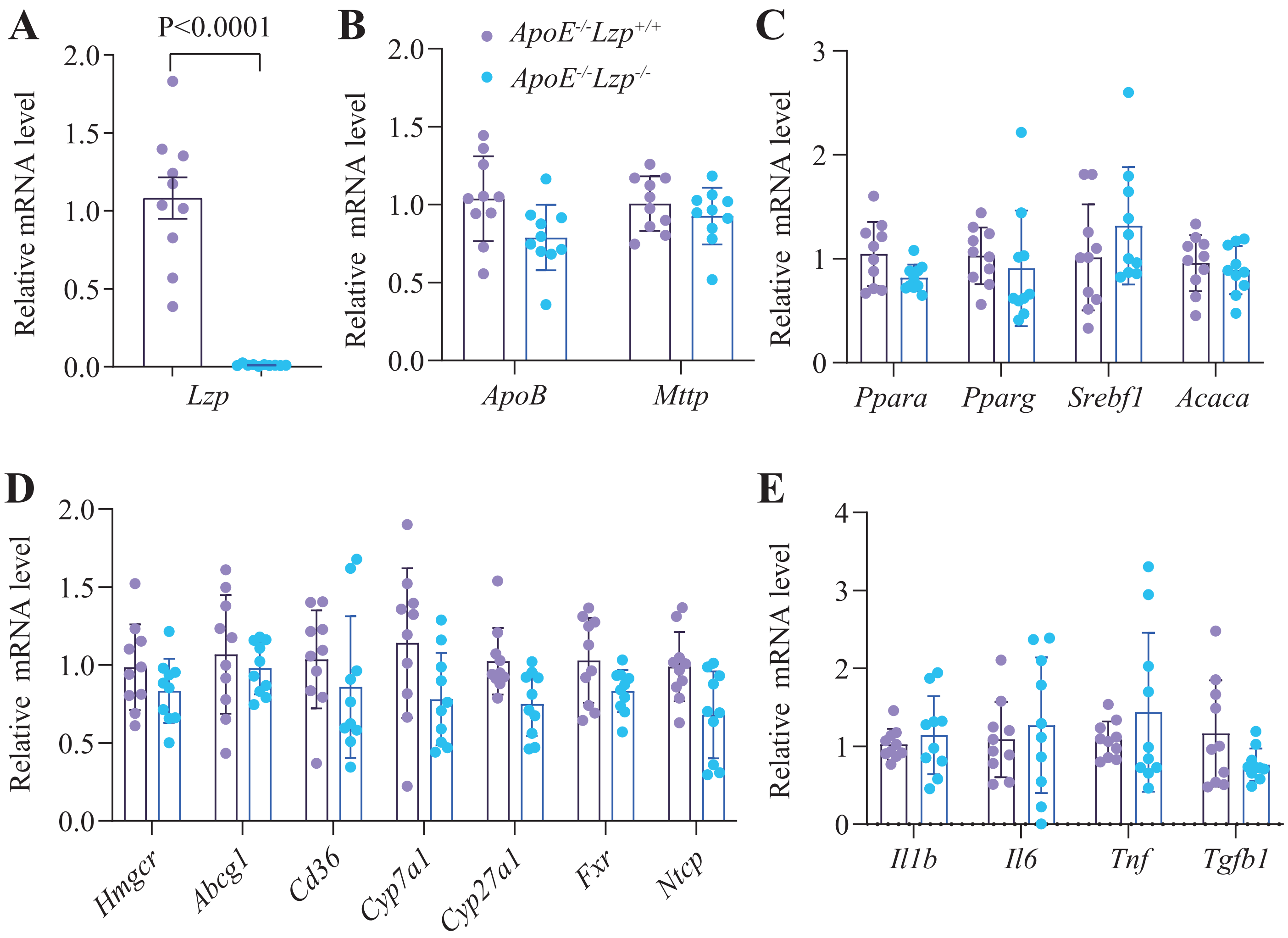
